# Elp1 function in placode-derived neurons is critical for proper trigeminal ganglion development

**DOI:** 10.1101/2024.07.12.603323

**Authors:** Margaret A. Hines, Lisa A. Taneyhill

## Abstract

**Background:** The trigeminal nerve is the largest cranial nerve and functions in somatosensation. Cell bodies of this nerve are positioned in the trigeminal ganglion, which arises from the coalescence of neural crest and placode cells. While this dual cellular origin has been known for decades, the molecular mechanisms controlling trigeminal ganglion development remain obscure. We performed RNAsequencing on the forming chick trigeminal ganglion and identified *Elongator acetyltransferase complex subunit 1* (*Elp1*) for further study. Mutations in *ELP1* cause familial dysautonomia (FD), a fatal disorder characterized by the presence of smaller trigeminal nerves and sensory deficits. While Elp1 has established roles in neurogenesis, its functions in placode cells during trigeminal gangliogenesis have not been investigated.

**Results:** To this end, we used morpholinos to deplete Elp1 from chick trigeminal placode cells. Elp1 knockdown decreased trigeminal ganglion size and led to aberrant innervation of the eye by placode-derived neurons. Trigeminal nerve branches exhibited fewer axons, and abnormal interactions between placode-derived neurons and neural crest cells were observed.

**Conclusions:** These findings reveal a new role for Elp1 in chick placode-derived neurons during trigeminal ganglion development. These results have potential high significance to provide new insights into trigeminal ganglion development and the etiology of FD.

**Bullet points:**

- Elp1 is expressed in undifferentiated neural crest cells and placode-derived neurons contributing to the trigeminal ganglion.
- Elp1 knockdown in trigeminal placode cells reduces trigeminal ganglion size.
- Elp1 depletion from trigeminal placode cells leads to aberrant target tissue innervation and disrupts proper neural crest-placodal neuron interactions in the trigeminal ganglion.

**Grant sponsor and number:** NIH R01DE024217 and NIH R03HD108480.

## INTRODUCTION

Cranial sensory nerves are components of the peripheral nervous system, located in the head, that are responsible for relaying sensory information to the central nervous system. The structure that houses the neuronal cell bodies and supporting glia of the sensory nerves is termed the ganglion. In the head, these include the trigeminal (V) and epibranchial (geniculate (facial VII), petrosal (glossopharyngeal IX) and nodose (vagal X)) ganglia. Nerves arising from these ganglia innervate diverse structures including the face, tongue, mouth, and digestive tract, and are derived from two embryonic cell populations, cranial neural crest cells and neurogenic placodes.

The trigeminal ganglion and its associated nerves are responsible for detecting pain, touch, and temperature sensations in the head and face (Koontz et al., 2023). Neural crest cells from the midbrain and rostral hindbrain regions, along with placode cells from the trigeminal placode, migrate and coalesce together to form this ganglion. While often referred to as one structure, the trigeminal placode is made up of two distinct placodes, the ophthalmic and maxillomandibular (Baker & Bronner-Fraser, 2001; D’Amico-Martel & Noden, 1983), each contributing to a specific lobe of the trigeminal ganglion. Unlike other cranial sensory ganglia, neurons derived from neural crest cells and placode cells are intermixed throughout this ganglion (D’Amico-Martel & Noden, 1983; Steventon et al., 2014). Reciprocal interactions between these cell types are required for proper trigeminal ganglion formation (D’Amico-Martel & Noden, 1983; Hamburger, 1961; Lwigale, 2001; Shiau & Bronner-Fraser, 2009); however, the molecules and pathways mediating ganglion development remain elusive.

Through RNAsequencing, we identified *Elongator acetyltransferase complex subunit 1* (*Elp1*) as a potential candidate for controlling chick trigeminal ganglion development. Mutations in human *ELP1* cause familial dysautonomia (FD), a neurodevelopmental and neurodegenerative disease. Evidence such as smaller trigeminal nerves in patients (and in FD mouse models), and symptoms such as decreased sensitivity to pain and temperature in the face, absent corneal reflexes, and reduced basal lacrimation, suggest the presence of trigeminal nerve deficits in FD (Jackson et al., 2014; Mass & Gadoth, 1994; Norcliffe-Kaufmann et al., 2017; Won et al., 2019). Studies have evaluated the function of Elp1 in neural crest cell populations using a conditional knockout mouse in which *Elp1* is deleted from neural crest cells and their derivatives through Wnt1-Cre-mediated recombination (George et al., 2013; Goffena et al., 2018, Jackson et al., 2014). Examination of trunk neural crest-derived sensory ganglia in this model revealed that Elp1 deletion caused second wave neuronal progenitors to differentiate early or undergo cell death (George et al., 2013), leading to fewer neurons and thus smaller ganglia.

Additional studies on Elp1 function in trunk sensory ganglia have been performed in the chick embryo (Abashidze et al., 2014; Hunnicutt et al., 2012). Depletion of Elp1 led to precocious neuronal differentiation, increased axon branching, and premature cell death in one study (Hunnicutt et al., 2012) while another identified abnormal axon branching, including an increase in branching points, neuron guidance abnormalities, and a role for Elp1 in axonal transport (Abashidze et al., 2014). These results add to the data from mouse models and strongly suggest roles for Elp1 in nerve outgrowth and/or target tissue innervation, neuron survival, and protein trafficking (George et al., 2013; Goffena et al., 2018; Li et al., 2020; Naftelberg et al., 2016, Jackson et al., 2014).

Using the conditional knockout mouse described above (George et al., 2013), we determined that loss of Elp1 in neural crest-derived neurons of the trigeminal ganglion causes axon outgrowth and target tissue innervation defects (Leonard et al., 2022). Understanding the full function of Elp1 in the trigeminal ganglion, however, requires investigating the role of Elp1 in placode-derived neurons as well. Since there is currently no Cre driver that only targets trigeminal placodes, we turned to the chick embryo. We first characterized Elp1 spatio-temporal distribution as the trigeminal ganglion initially assembles from undifferentiated neural crest cells and placode-derived neurons. Our data show that Elp1 is expressed in migratory cranial neural crest cells and later in undifferentiated neural crest cells and placode-derived neurons contributing to the trigeminal ganglion. Next, we performed morpholino-mediated knockdown of Elp1 in trigeminal placode cells and uncovered a sustained negative impact on trigeminal ganglion development. Our results are the first to describe Elp1 expression in the forming chick trigeminal ganglion and point to critical functions for Elp1 in placode-derived neurons during trigeminal ganglion development, providing additional insight into the etiology of trigeminal nerve deficits in FD.

## RESULTS

### Elp1 is dynamically expressed throughout trigeminal ganglion development

Characterization of Elp1 protein expression was performed in stages, with embryos grouped every 12-24 hours, as this correlates with the timeframe of major developmental events (e.g., neural crest cell migration, initial intermixing of neural crest cells and placode-derived neurons, and condensation of neural crest cells and placode-derived neurons). A control experiment was first performed to ascertain the specificity of the Elp1 antibody (Figure 1). Immunostaining for Elp1 (Figure 1A,B) was performed in combination with Sox10 (Figure 1A,C) and Tubb3 (Figure 1A,D) to identify neural crest cells and placode-derived neurons of the trigeminal ganglion, respectively (Moody et al., 1989). Next, immunostaining was conducted in the absence of Elp1 antibody but in the presence of a secondary antibody specific for Elp1 (Figure 1E,F), accompanied by the same combination of Sox10 (Figure 1E,G) and Tubb3 (Figure 1E,H) antibodies. This latter experiment showed no signal for Elp1, confirming specificity of the Elp1 antibody, which was then used for subsequent characterization of the spatio-temporal expression pattern of Elp1.

**Figure 1:**
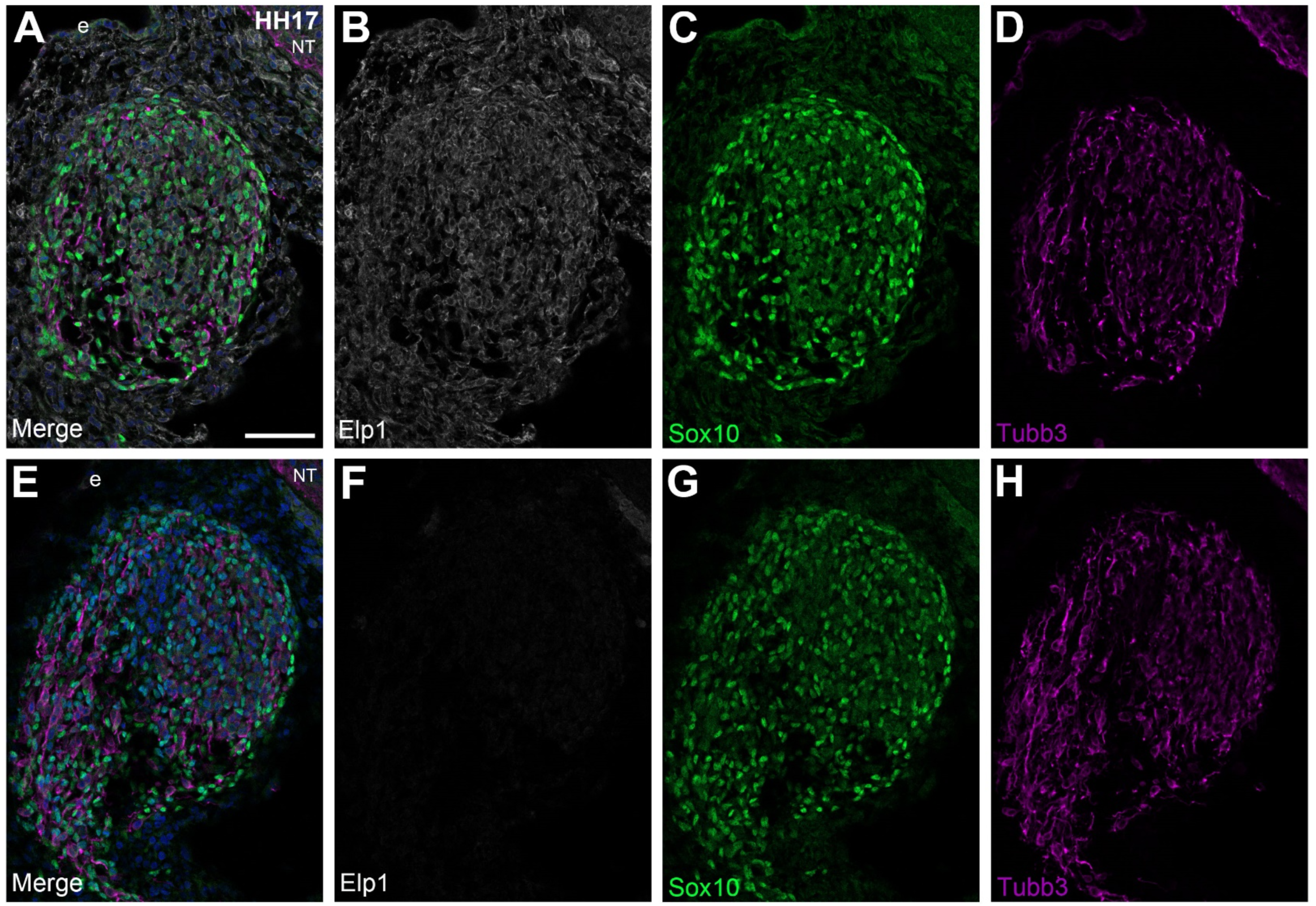
Secondary antibody only control for Elp1 immunohistochemistry supports specificity of the Elp1 antibody. Representative transverse section through the forming trigeminal ganglion ophthalmic lobe (E2.5, HH17, n = 3) followed by immunohistochemistry for Elp1 (A,B, white), Sox10 (A,C,E,G, green, labels neural crest cells), and Tubb3 (A,D,E,H, purple, labels placode-derived neurons) with corresponding merged images of all channels with DAPI (A,E, blue, marks all nuclei). Secondary antibody only control with absence of primary antibody for Elp1 (E,F, white) yields no detectable Elp1 fluorescence (E,F). Scale bar in (A) is 50μm and applies to all images. Abbreviations: e = ectoderm; NT = neural tube.

The first stage group investigated was embryonic day (E) 1.5 (Hamburger-Hamilton (HH)11/12), when neural crest cells are migrating away from the dorsal neural tube to their respective locations. At these timepoints, placode cells reside in the ectoderm and have yet to start their delamination and migration. Within Sox10-positive migratory neural crest cells, Elp1 expression is punctate in appearance (Figure 2B-D, arrowheads). Furthermore, Elp1 is observed in the ectoderm containing the placodal precursors, demonstrated by its presence in E-Cadherin-positive cells. This ectodermal expression appeared to be concentrated at the apical side of the cells (Figure 2B,C,E, arrows).

**Figure 2:**
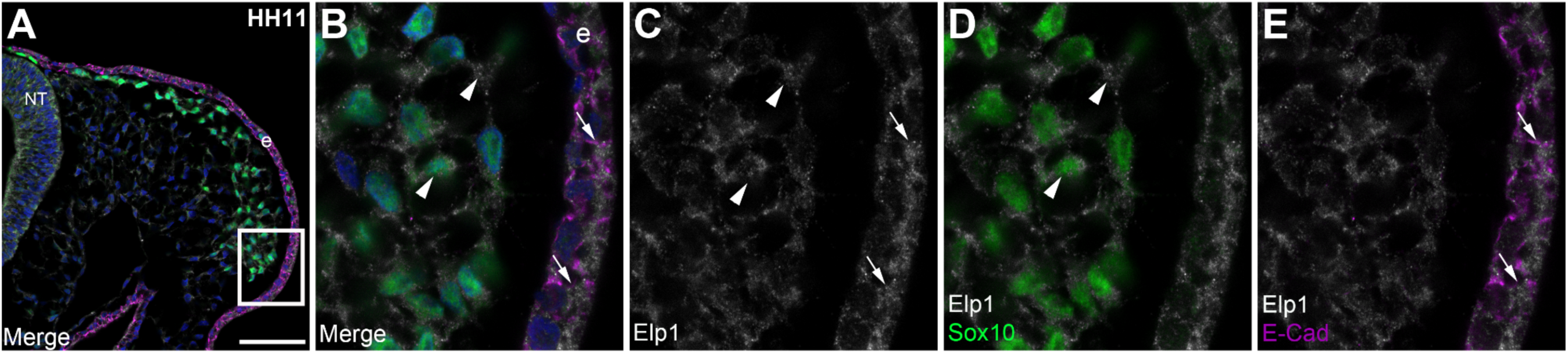
Elp1 is expressed in migratory neural crest cells and the surface ectoderm containing trigeminal placode cells prior to trigeminal ganglion assembly. Representative transverse section through an E1.5 (HH11, n = 3) embryo midbrain followed by immunohistochemistry for Elp1 (A-E, white), Sox10 (A,B,D, green, labels migratory neural crest cells), and E-Cadherin (E-Cad, A,B,E, purple, labels the ectoderm), with corresponding merged images of all channels with DAPI (A,B, blue, marks all nuclei). A higher magnification image of the boxed region in (A) is shown in (B-E), with merge images of Elp1 and Sox10 (D, white and green, respectively) and Elp1 and E-Cad (E, white and purple, respectively). Arrowheads point to Elp1 in migratory neural crest cells (B-D), while arrows indicate Elp1 in the ectoderm (B,C,E). Scale bar in (A) is 50μm and applies to all images but is 10μm for (B-E). Abbreviations: NT = neural tube; e = ectoderm.

At E2 (HH13-15), placode cells start to delaminate from the ectoderm and differentiate into neurons, identified by their immunoreactivity for Tubb3 (Figure 3A,B,E). While Tubb3 eventually labels all neurons in the trigeminal ganglion, these neurons are placode-derived at all stages in this expression profile, since neural crest cells differentiate much later (D’Amico-Martel & Noden, 1980). Elp1 expression (Figure 3A-E) is still present in the ectoderm (Figure 3C); however, the apical distribution of Elp1 has changed, with enrichment reduced compared to the previous stage group. We found continued and dynamic punctate expression of Elp1 in the cytoplasm and nuclei (Figure 3B-D, arrowheads) of Sox10-positive neural crest cells. Additionally, at these stages, Elp1 expression is noted in the cytoplasm of Tubb3-positive neurons (Figure 3B,C,E, arrows).

**Figure 3:**
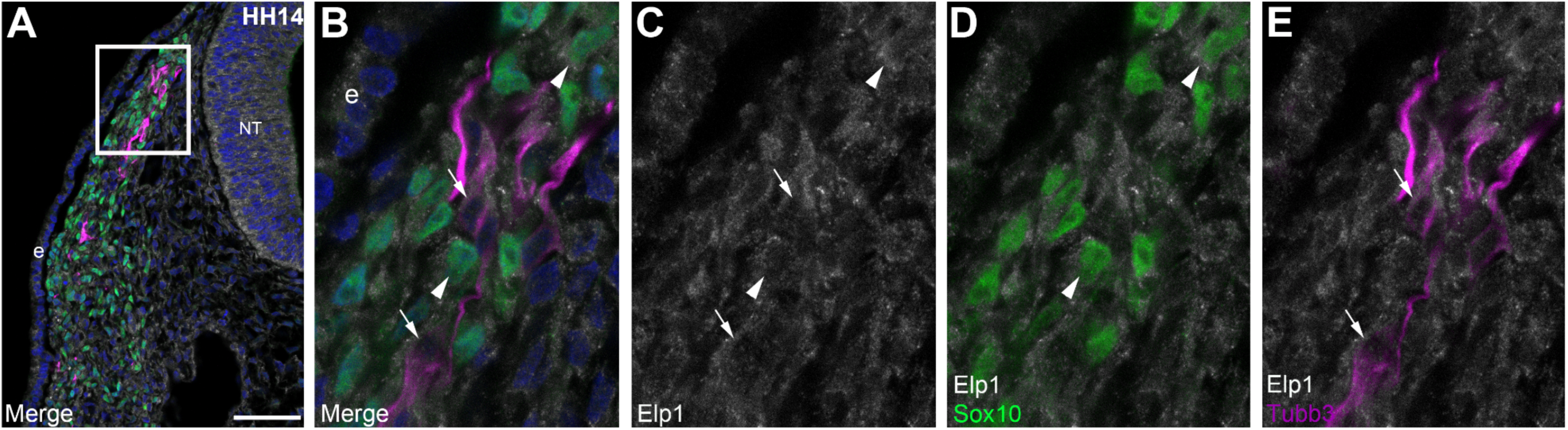
Elp1 is expressed in neural crest cells and placode-derived neurons of the trigeminal ganglion during early placode cell delamination and migration. Representative transverse section through the forming trigeminal ganglion ophthalmic lobe (E2, HH14, n = 3) followed by immunohistochemistry for Elp1 (A-E, white), Sox10 (A,B,D, green, labels neural crest cells), and Tubb3 (A,B,E, purple, labels placode-derived neurons), with corresponding merged images of all channels with DAPI (A,B, blue, marks all nuclei). A higher magnification image of the boxed region in (A) is shown in (B-E), with merge images of Elp1 and Sox10 (D, white and green, respectively) and Elp1 and Tubb3 (E, white and purple, respectively). Arrowheads point to Elp1 in migratory neural crest cells (B-D), while arrows indicate Elp1 in placode-derived neurons (B,C,E). Scale bar in (A) is 50μm and applies to all images but is 10μm for (B-E). Abbreviations: e = ectoderm.

At E2.5 (HH16-17), neural crest cells and placode-derived neurons have coalesced to form the trigeminal ganglion proper, where a more defined “teardrop” or “semilunar” shape is then observed in sections. Expression of Elp1 in Sox10-positive neural crest cells (Figure 4A-D, arrowheads) and the cytoplasm of Tubb3-positive neurons (Figure 4A,B,C,E, arrows) remains the same as in the previous stage grouping. Lastly, during E3-3.5 (HH18-20), a more defined trigeminal ganglion structure is present, with more placode-derived neurons surrounded by undifferentiated neural crest cells. Elp1 expression (Figure 5A-E, white) remains the same as observed in the previous stage groups, with ectodermal (Figure 5A), neural crest cell (Figure 5A-D, arrowheads), and neuronal (Figure 5A,B,C,E, arrows) expression noted. Taken together, these results indicate that Elp1 is expressed in the proper cell types and at the correct time to function in trigeminal ganglion development.

**Figure 4:**
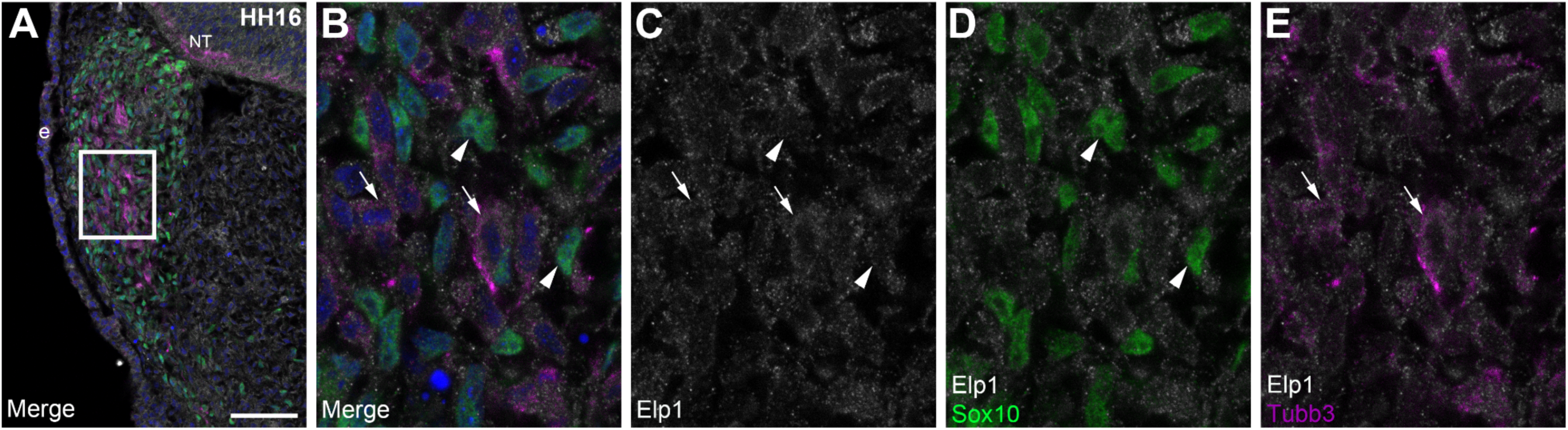
Elp1 is expressed in neural crest cells and placode-derived neurons of the trigeminal ganglion during initial coalescence and condensation. Representative transverse section through the forming trigeminal ganglion ophthalmic lobe (E2.5, HH16, n = 3) followed by immunohistochemistry for Elp1 (A-E, white), Sox10 (A,B,D, green, labels neural crest cells), and Tubb3 (A,B,E, purple, labels placode-derived neurons), with corresponding merged images of all channels with DAPI (A,B, blue, marks all nuclei). A higher magnification image of the boxed region in (A) is shown in (B-E), with merge images of Elp1 and Sox10 (D, white and green, respectively) and Elp1 and Tubb3 (E, white and purple, respectively). Arrowheads point to Elp1 in migratory neural crest cells (B-D), while arrows indicate Elp1 in placode-derived neurons (B,C,E). Scale bar in (A) is 50μm and applies to all images but is 10μm for (B-E). Abbreviations: e = ectoderm; NT = neural tube.

**Figure 5:**
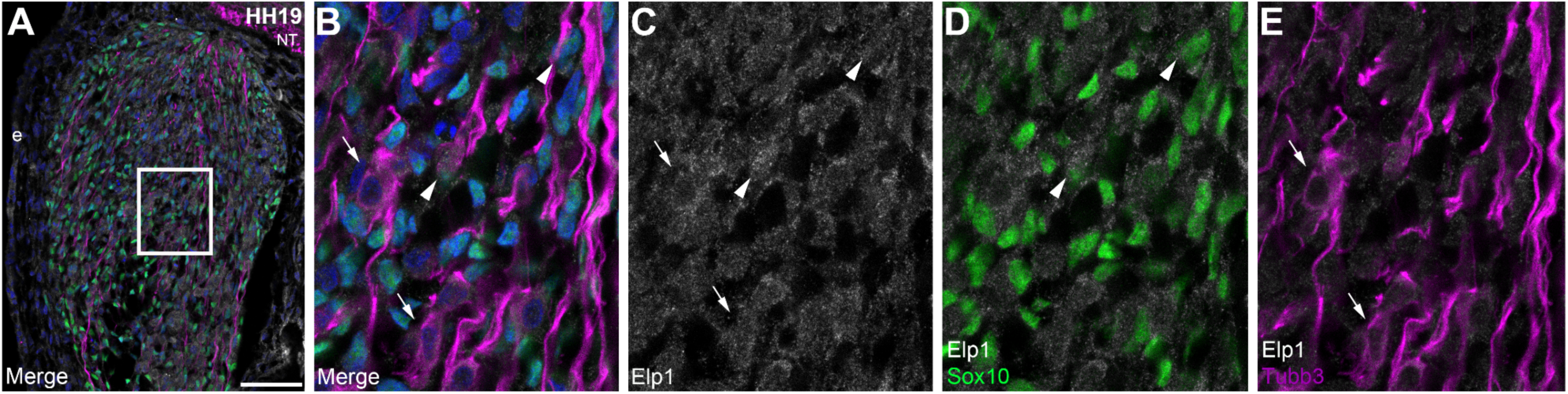
Elp1 expression persists in neural crest cells and placode-derived neurons of the trigeminal ganglion at later stages. Representative transverse section through the forming trigeminal ganglion ophthalmic lobe (E3, HH19, n = 3) followed by immunohistochemistry for Elp1 (A-E, white), Sox10 (A,B,D green, labels neural crest cells), and Tubb3 (A,B,E, purple, labels placode-derived neurons), with corresponding merged images of all channels with DAPI (A,B, blue, marks all nuclei). A higher magnification image of the boxed region in (A) is shown in (B-E), with merge images of Elp1 and Sox10 (D, white and green, respectively) and Elp1 and Tubb3 (E, white and purple, respectively). Arrowheads point to Elp1 in migratory neural crest cells (B-D), while arrows indicate Elp1 in placode-derived neurons (B,C,E). Scale bar in (A) is 50μm and applies to all images but is 10μm for (B-E). Abbreviations: e = ectoderm.

### Elp1 morpholinos are effective at reducing Elp1 protein levels in trigeminal placode cells

A 1:1 mixture of an *Elp1* translation-blocking morpholino and *Elp1* splice-blocking morpholino (referred to hereafter as MOs), or a standard Control MO (GeneTools, LLC), was electroporated at E1 (HH10-11) using a unilateral ectodermal electroporation method (Shah et al., 2017; Shiau et al., 2008) to target trigeminal placode cells prior to delamination. Immunoblotting was then performed with an Elp1 antibody previously validated for this method (Li et al., 2020). This allowed us to visualize and measure the level of Elp1 reduction to confirm efficacy of MOs. Immunoblotting revealed six bands immunoreactive to the Elp1 antibody, five of which showed varying levels of knockdown ranging from 14-56% (Figure 6). The band at 150 kDa, however, showed no reduction in intensity after introduction of the Elp1 MOs. These findings indicate that this combination of Elp1 MOs can deplete Elp1 protein levels, providing a means by which to evaluate Elp1 function in the trigeminal ganglion.

**Figure 6:**
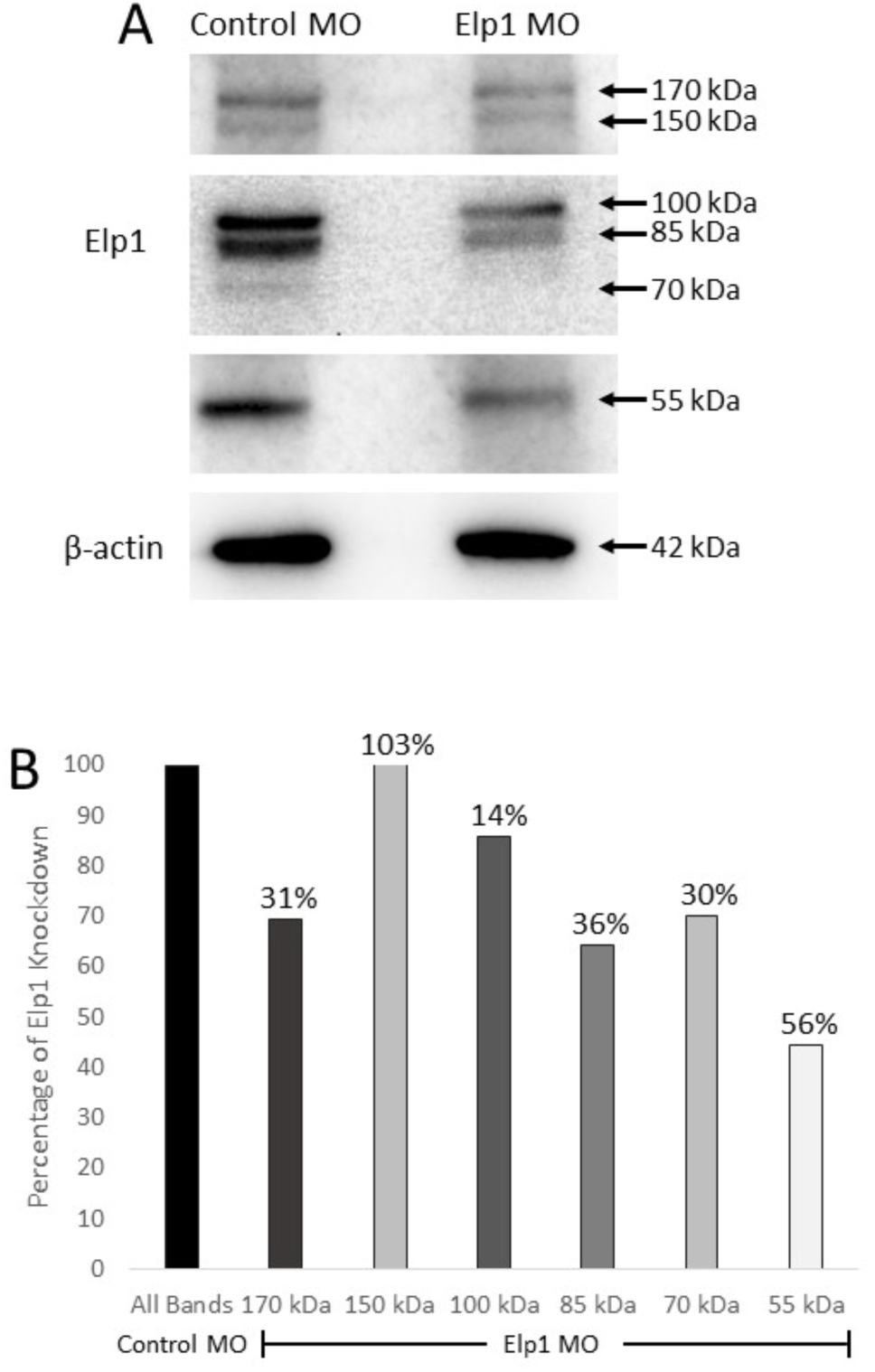
Elp1 MOs reduce Elp1 protein levels in the trigeminal ganglion. The knockdown efficiency of a 1:1 ratio of electroporated Elp1 translation-blocking and splice-blocking MOs was assessed using immunoblotting on whole-cell lysates prepared from dissected Elp1 MO- or Control MO-positive trigeminal ganglia. (A). Six bands were immunoreactive to the Elp1 antibody and when normalized to β-actin, with five showing reduced intensities in the Elp1 MO-treated tissue compared to the Control MO-treated tissue (B).

### Elp1 knockdown in trigeminal placode cells decreases the area of the trigeminal ganglion during early stages of trigeminal ganglion assembly

To investigate a role for Elp1 in trigeminal ganglion development, Elp1 MOs were electroporated into trigeminal placode cells as described above. Trigeminal ganglion development was first evaluated at ∼E2.5 (HH15-17), when placode-derived neurons and migratory neural crest cells are intermingling, through whole-mount immunohistochemical staining for Tubb3 to label placode-derived neurons. Strikingly, defects were already apparent at these early stages of trigeminal ganglion development (Figure 7). On the contralateral (control) side of the embryo, the trigeminal ganglion formed normally, with ophthalmic and maxillomandibular lobes starting to become defined (Figure 7A, arrowhead, arrow, respectively). On the Elp1 MO-treated side, knockdown of Elp1 negatively impacted the development of the ophthalmic branch of the trigeminal ganglion, with disorganization of neurons apparent (Figure 7B). Measurement of the area occupied by experimental and contralateral control side trigeminal ganglia revealed a statistically significant decrease upon Elp1 knockdown compared to control (Figure 7C, p = 0.0014). Importantly, electroporation of a scrambled Control MO into trigeminal placode cells led to no phenotypic effect on trigeminal ganglion development, with results comparable to those observed on the contralateral sides of both Elp1 MO- and Control MO-treated embryos (Figure 8). Therefore, the contralateral control (nonelectroporated) sides of Elp1 MO-treated embryos were used as subsequent controls in future experiments to account for any potential variation that normally occurs during embryonic development. Altogether, these data reveal early effects on trigeminal ganglion development upon depletion of Elp1 from placode cells.

**Figure 7:**
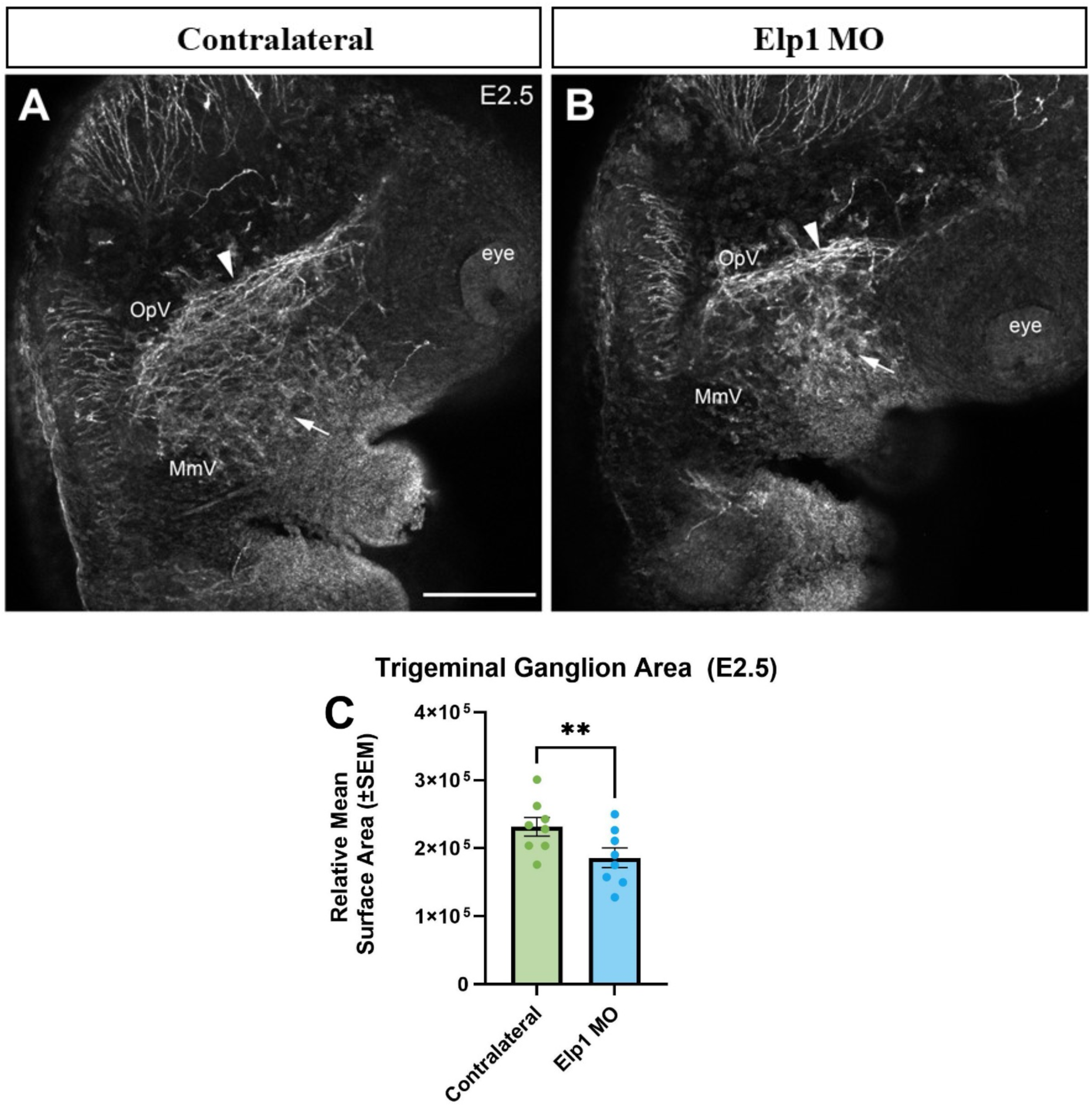
Reduction of Elp1 in trigeminal placode cells decreases trigeminal ganglion area and impacts neuronal organization. Representative max intensity projections of confocal Z-stacks showing the contralateral control (A) or Elp1 MO-treated (B) trigeminal ganglion at E2.5 (HH16) after Tubb3 whole-mount immunohistochemistry (white). Arrowheads point to disorganization of ophthalmic branch projections, while arrows show the forming maxillomandibular branch. Scale bar in (A) is 250µm and applies to (B). (C) Quantification of the area of contralateral (green, n = 7) and Elp1 MO-treated (blue, n = 7) trigeminal ganglia demonstrating statistical significance (p = 0.0014, paired t-test). Abbreviations: OpV = ophthalmic lobe; MmV = maxillomandibular lobe.

**Figure 8:**
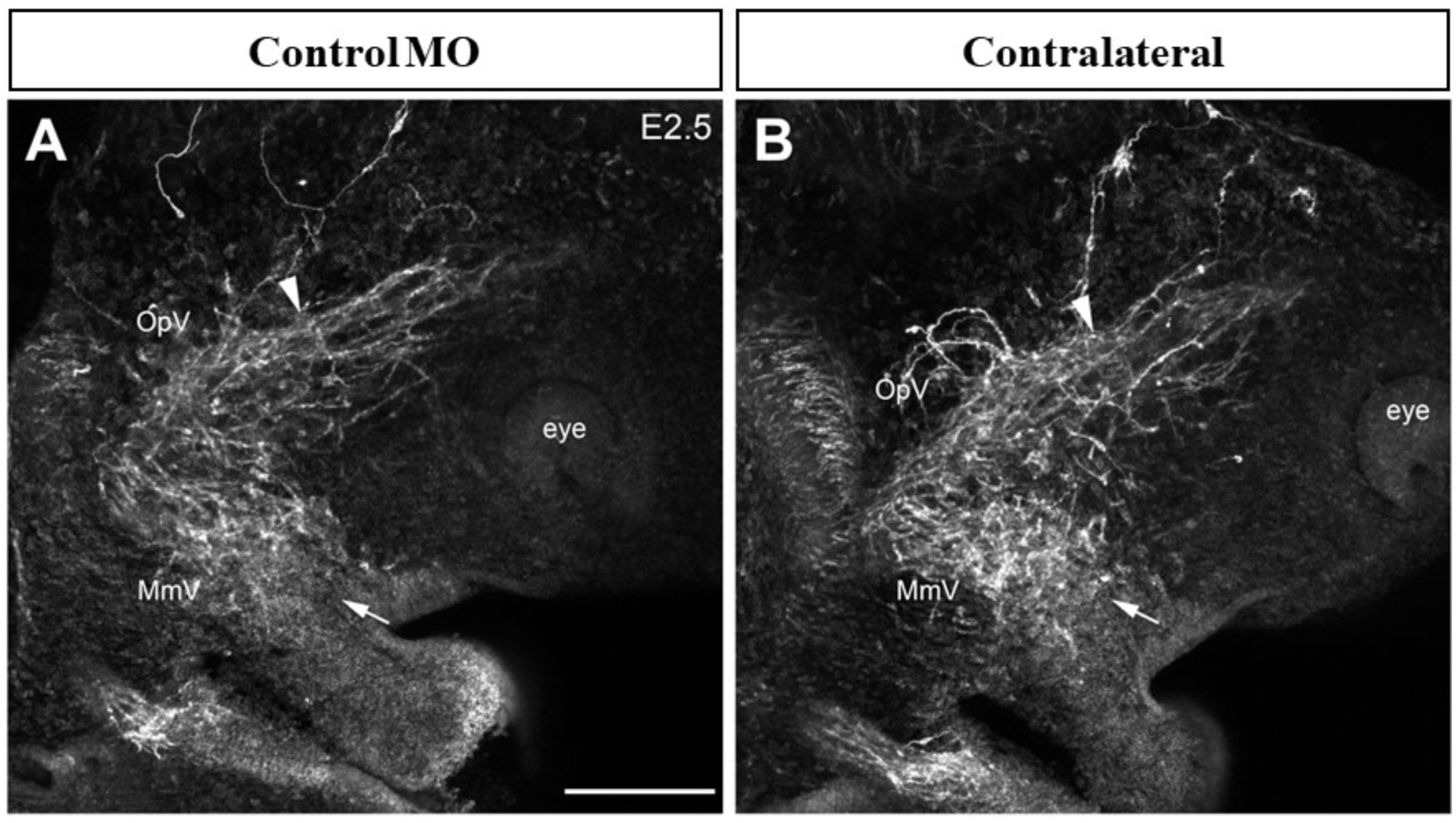
Electroporation of the scrambled Control MO into trigeminal placode cells does not alter trigeminal ganglion development. Representative max intensity projections of confocal Z-stacks showing the Control MO-treated (A) or contralateral control (B) trigeminal ganglion at E2.5 (HH16, n = 3) after Tubb3 whole-mount immunohistochemistry (white). Arrowheads point to normal ophthalmic branch projections, while arrows show a properly forming maxillomandibular branch. Scale bar in (A) is 250µm and applies to (B). Abbreviations: OpV = ophthalmic lobe; MmV = maxillomandibular lobe.

### Elp1 knockdown in trigeminal placode cells causes persistent reduction in trigeminal ganglion area and negatively affects innervation of the eye

One day later, extensive development of the trigeminal ganglion has occurred through further condensation of neural crest cells and placode-derived neurons (∼E3.5, HH19-20). Immunohistochemical staining for Tubb3 (marking placode-derived neurons) revealed distinct ophthalmic, maxillary, and mandibular branches on the contralateral control trigeminal ganglion (Figure 9A, arrowhead, arrows, respectively). After Elp1 knockdown in trigeminal placode cells (Figure 9B), the area encompassed by the trigeminal ganglion remained significantly reduced (Figure 9C, p = 0.0425). Furthermore, evaluation of the forming maxillary and mandibular branches indicated a dispersal of axonal projections (Figure 9B, arrows) compared to the control. However, analysis of the widths for each branch revealed no significant changes (data not shown).

**Figure 9:**
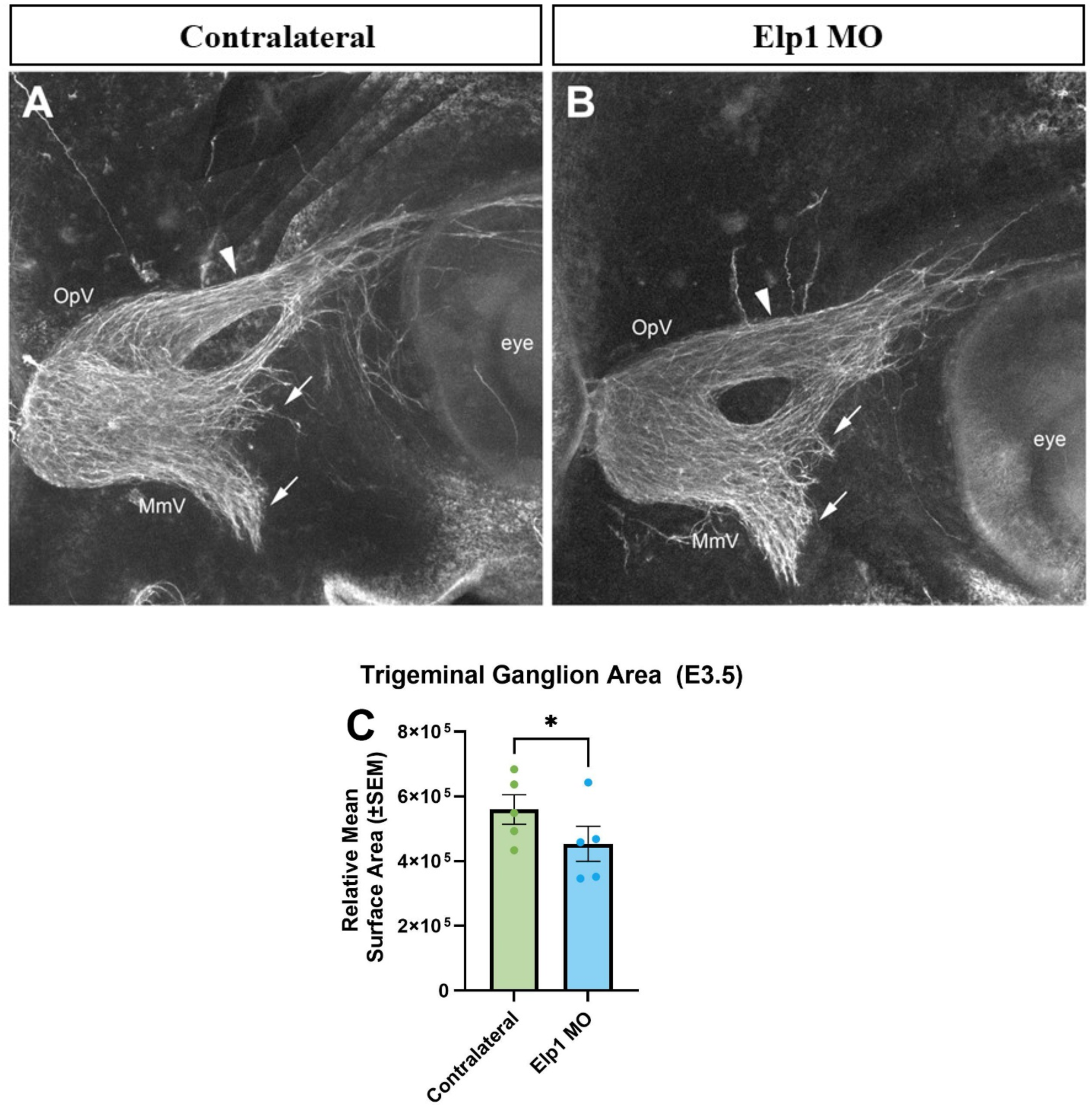
Elp1 depletion reduces the area of the trigeminal ganglion at later developmental stages. Representative max intensity projections of confocal Z-stacks showing the contralateral control (A) or Elp1 MO-treated (B) trigeminal ganglion at E3.5 (HH19-20) after Tubb3 whole-mount immunohistochemistry (white). Arrowheads point to defasciculated and/or dispersed ophthalmic branch axons, while arrows show dispersion of maxillomandibular branch axons. Scale bar in (A) is 250µm and applies to (B). (C) Quantification of the area of contralateral (green, n = 7) and Elp1 MO-treated (blue, n = 5) trigeminal ganglia demonstrating statistical significance (p = 0.0425, paired t-test). Abbreviations: OpV = ophthalmic lobe; MmV = maxillomandibular lobe.

Further examination of the ophthalmic branch innervation pattern showed normal axonal branching and projections targeting the contralateral control eye (Figure 10A, arrowhead). Innervation of the Elp1 MO-treated side eye, however, was abnormal, with aberrant axonal branching and projections (Figure 10B, arrowhead). Quantification of this region (i.e., area occupied by ophthalmic axons) revealed a statistically significant reduction in the innervation of the eye after Elp1 knockdown (Figure 10C, p = 0.004) compared to the contralateral control side. Collectively, these data point to long term defects in the outgrowth of axons from placode-derived neurons after Elp1 knockdown.

**Figure 10:**
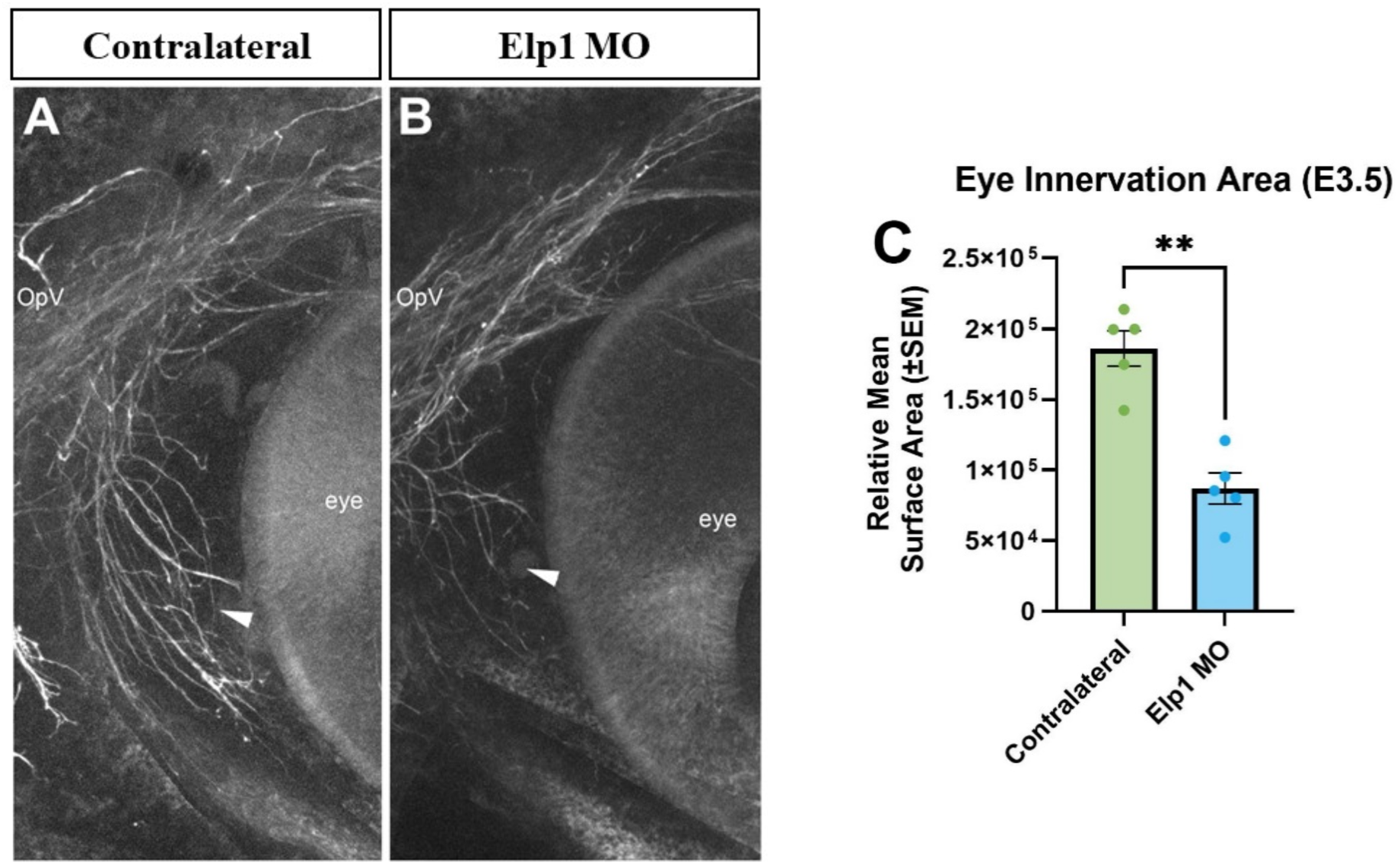
Reduction of eye innervation is observed after Elp1 knockdown. Representative max intensity projections of confocal Z-stacks showing the contralateral control (A) or Elp1 MO-treated (B) trigeminal ganglion at E3.5 (HH19-20) after Tubb3 whole-mount immunohistochemistry (white) with a focus on the innervation of the eye by the ophthalmic branch. Arrowheads point to axonal projections innervating the eye. Scale bar in (A) is 300µm and applies to (B). (C) Quantification of the area of contralateral (green, n = 5) and Elp1 MO-treated (blue, n = 5) ophthalmic innervation of the eye demonstrating statistical significance (p = 0.004, paired t-test). Abbreviations: OpV = ophthalmic lobe.

### Elp1 knockdown in trigeminal placode leads to disorganization of placodal neurons

To better discern potential changes at the cellular level after Elp1 knockdown, section immunohistochemistry was performed to identify placode-derived neurons and undifferentiated neural crest cells. At early stages of development (E2.5, HH15-17), the contralateral control trigeminal ganglion (Figure 11A-D) possessed the typical organization of Sox10-positive neural crest cells (Figure 11A,C, arrowheads) surrounding Tubb3-positive placode-derived neurons (Figure 11A,D, arrows). Evaluation of the Elp1 MO-treated trigeminal ganglion (Figure 11E-H), however, uncovered a change in the arrangement of neural crest cells and placode-derived neurons after Elp1 knockdown. Condensation of placode-derived neurons was negatively impacted, with Tubb3 immunoreactive cells more dispersed (Figure 11E,H, arrows). Additionally, neuronal projections were smaller or even absent, and placode-derived neurons appeared disorganized and more dispersed. Furthermore, Sox10-positive neural crest cells appeared less condensed (Figure 11E,G, arrowheads). Taken together, these findings suggest that knockdown of Elp1 affects interactions between neural crest cells and placode-derived neurons, including as these neurons begin to extend and bundle axons.

**Figure 11:**
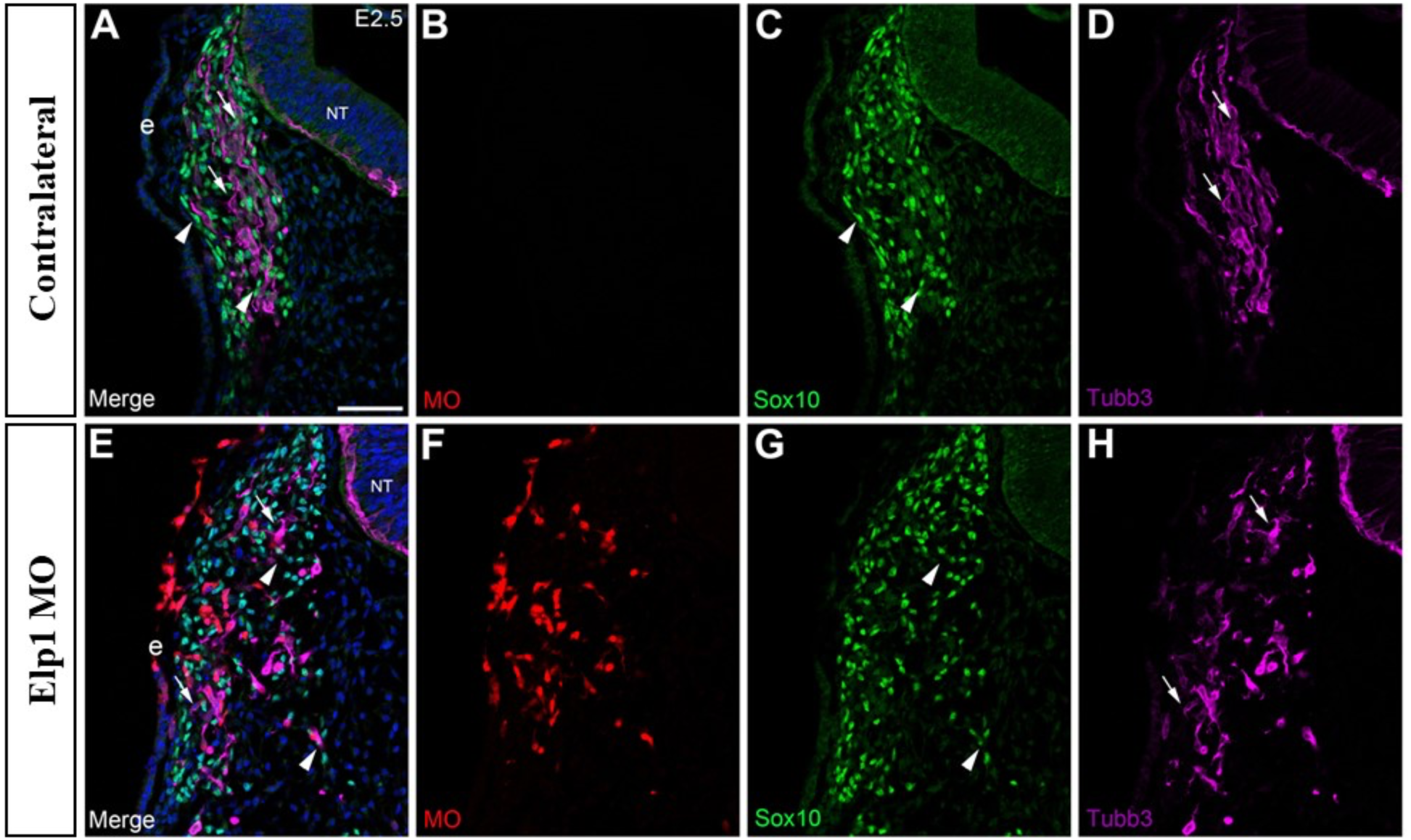
Elp1 knockdown increases cell dispersal within the trigeminal ganglion. Representative transverse section through the forming ophthalmic lobe of the trigeminal ganglion of an Elp1 MO-treated embryo at E2.5 (HH16), with the contralateral control and Elp1 MO-treated sides shown in (A-D) and (E-H), respectively. Immunohistochemistry for Sox10 (C,G, green, labels neural crest cells) and Tubb3 (D,H, purple, labels placode-derived neurons) was conducted on tissue sections. MO-positive cells are visualized by the lissamine tag on the MO, which fluoresces red (F; none in B), with corresponding merged images of all channels with DAPI (A,E, blue, marks all nuclei). Arrowheads denote Sox10-positive neural crest cells (A,C,E,G), while arrows point to Tubb3-positive neurons (A,D,E,H). Scale bar in (A) is 50µm and applies to all images. Abbreviations: NT = neural tube; e = ectoderm.

### Deficits in trigeminal ganglion body size and neuronal projections are apparent at later developmental timepoints after Elp1 knockdown

Later in trigeminal ganglion development (E3.5, HH19-20), neural crest cells and placode-derived neurons are tightly condensed, with neurons extending axons to target tissues. At this stage, the ophthalmic projections are easily distinguishable in cross-sections through the trigeminal ganglion (Figure 12A,E). The projections from the contralateral control side trigeminal ganglion exhibited well-organized Tubb3-positive axons (Figure 12A,D, arrow), with Sox10-positive neural crest cells lining the axons as expected (Figure 12A,C, arrowhead). After Elp1 knockdown (Figure 12E-H), axon projections identified by Tubb3 immunoreactivity (Figure 12E,H, arrow) were abnormal, with a large degree of disorganization and dispersal, as seen at earlier stages (Figure 11). Moreover, the organization of Sox10-positive neural crest cells along some of the neuronal projections was also aberrant (Figure 12E,G, arrowhead). Qualitatively, there is an apparent reduction in neural crest cell number compared to the contralateral control side. These results further support a role for Elp1 in controlling the outgrowth and fasciculation of axons from placode-derived neurons during trigeminal ganglion development.

**Figure 12:**
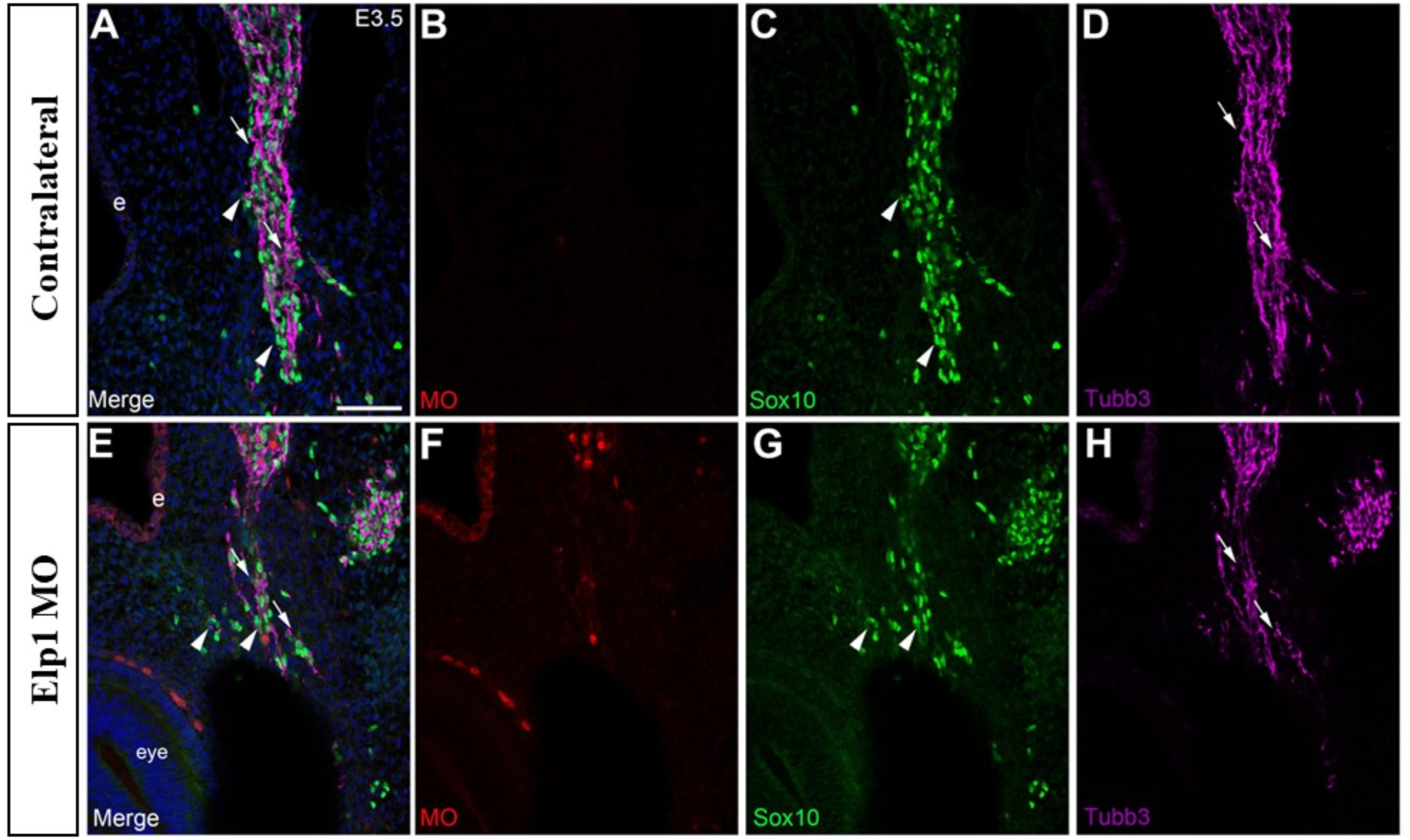
Abnormal outgrowth of neuronal projections and disorganization of neural crest cells is noted after Elp1 knockdown. Representative transverse section through the forming ophthalmic branch of the trigeminal ganglion of an Elp1 MO-treated embryo at E3.5 (HH20), with the contralateral control (A-D) and Elp1 MO-treated (E-H) sides shown. Immunohistochemistry for Sox10 (A,C,E,G, green, labels neural crest cells) and Tubb3 (A,D,E, H, purple, labels placode-derived neurons). MO-positive cells are visualized by the lissamine tag on the MO, which fluoresces red (F; none in B), with corresponding merged images of all channels with DAPI (A,E,blue, marks all nuclei). The arrowhead indicates Sox10-positive neural crest cells (A,C,E,G), while the arrow marks Tubb3-positive neurons (A,D,E,H). Scale bar in (A) is 50µm and applies to all images. Abbreviations: e = ectoderm.

## DISCUSSION

Development of the cranial sensory ganglia is a very complex and highly coordinated process. For example, neural crest cells and placode-derived neurons must work together through reciprocal interactions to form the trigeminal ganglion (D’Amico-Martel & Noden, 1983; Hamburger, 1961; Lwigale, 2001; Shiau & Bronner-Fraser, 2009), which is involved in the perception of many sensations in the head and face, including pain, touch, and temperature (Koontz et al., 2023). While the cellular origin of the trigeminal ganglion is firmly established, the molecules involved in trigeminal ganglion development are still not known. To fill this knowledge gap, we identified Elp1 as a candidate for regulating chick trigeminal gangliogenesis. Intriguingly, mutations in *ELP1* cause FD, a fatal disorder that broadly affects the nervous system. Symptoms associated with FD, along with prior studies in mouse models, reveal trigeminal nerve developmental defects. While the expression and function of Elp1 in neural crest-derived trigeminal neurons has been characterized in a mouse model of FD, the role of Elp1 in trigeminal placode cells had yet to be studied. To this end, we used the chick embryo to examine Elp1 expression and function in trigeminal placode cells and subsequent trigeminal ganglion development. The results of these experiments showed that knockdown of Elp1 in trigeminal placode cells leads to disruption of trigeminal ganglion development in specific ways.

### Elp1 is expressed in the proper spatio-temporal pattern to function in trigeminal ganglion development

While our data revealed ubiquitous Elp1 expression in the chick embryo, we observed enrichment in the ectoderm, including where the trigeminal placodes will form, migratory neural crest cells, and neural crest cells and placode-derived neurons within the trigeminal ganglion. Section immunohistochemistry revealed a punctate cellular distribution of Elp1 throughout the timepoints characterized (E1.5-3.5, HH11-20). Interestingly, previous studies in the chick trunk showed that Elp1 was not expressed in migratory neural crest cells but was detected in the cytoplasm and axons of neural crest-derived dorsal root ganglion neurons (Hunnicutt et al., 2012). A later study also reported Elp1 expression only in post-mitotic neurons of the dorsal root ganglia (Abashidze et al., 2014). These immunohistochemistry data, however, were acquired using a different Elp1 antibody that is no longer commercially available, with a different epitope sequence than the antibody used in our studies. Therefore, expression differences may be due to epitope location and/or antibody affinities. In addition to the antibodies used, differences between trunk and cranial development are relatively common (D’Amico-Martel & Noden, 1980) and may explain the expression pattern and distribution for Elp1 uncovered in our current studies.

In the E10.5 mouse trigeminal ganglion, Elp1 is enriched in the cytoplasm and axons of neurons, with little to no Elp1 present in undifferentiated neural crest cells and a general lack of nuclear staining observed (Leonard et al., 2022). While these immunohistochemical results are consistent with the expression pattern we observed in the forming chick trigeminal ganglion, expression differences could be attributed to the stages examined. At E10.5 in the mouse, neural crest cells are poised to begin differentiating into trigeminal neurons, beginning at E11 (Karpinski et al., 2016). The latest stages examined in our chick studies, however, are still at least a day before neural crest cells start differentiating into neurons. Importantly, Elp1 expression has yet to be examined at earlier developmental timepoints in the mouse when neural crest cells begin their migration. Lastly, our chick data revealed punctate expression of Elp1 in the cell types in which Elp1 was expressed. Given the known localization of Elp1 to intracellular vesicles (Li et al., 2020), immunohistochemistry with additional antibodies to label these vesicles will aid in identifying the subcellular location of Elp1.

### Elp1 knockdown in placode cells negatively impacts trigeminal ganglion development

To evaluate Elp1 function, we used a combination of splice- and translation-blocking MOs to deplete Elp1 from trigeminal placode cells. Knockdown was verified through immunoblotting, which revealed decreases in five bands that were Elp1-immunoreactive except for a band at 150 kDa (assumed to be a background band since its levels did not change). Multiple bands at different molecular weights could point to various Elp1 isoforms, differential modification of Elp1, and/or proteolytic processing of Elp1 that occurs throughout embryonic development, particularly since immunoblotting was performed on tissue dissected from pooled embryos at E3.5 (HH19-20), which encompasses two embryonic stages.

Using these MOs, we found a reduction in the area occupied by the trigeminal ganglion during initial formation at E2.5 (HH15-17). Examination of sections through the trigeminal ganglion at this stage revealed that, in some embryos, placode cells may be remaining in the ectoderm. If these placode cells do not delaminate, they may eventually undergo apoptosis, preventing them from contributing to the trigeminal ganglion. This would be in keeping with previous studies investigating Elp1 in trunk neural crest-derived sensory neurons, where increased cell death was also seen (Abashidze et al., 2014; George et al., 2013; Hunnicutt et al., 2012; Leonard et al., 2022), and could be further verified by TUNEL staining. Additionally, both placode-derived neurons and neural crest cells appeared more dispersed, suggesting the reciprocal interactions between these two populations have been abrogated. This non-cell autonomous effect has been previously observed after knockdown of gene expression in one trigeminal ganglion precursor cell type, leading to defects associated with the other precursor cell type (Shiau & Bronner-Fraser, 2009; Wu & Taneyhill, 2019; Wu et al., 2014; Halmi et al., 2024). Counting of labeled of placode-derived neurons and undifferentiated neural crest cells on tissue sections will allow these phenotypes to be quantified. A day later in development (E3.5, HH19-20), the decrease in area occupied by the trigeminal ganglion persisted, and we observed a statistically significant reduction in the ophthalmic branches innervating the eye. Abnormal outgrowth visualized in section further supports defects in axonal development, showing disorganization of axon projections. The distribution of some neural crest cells is also aberrant, consistent with observations at E2.5.

Previous studies have evaluated Elp1 function in the chick embryo but in the neural crest cell-derived trunk dorsal root ganglia. One report found Elp1-depleted trunk neural crest cells to undergo proper induction, delamination, and migration; however, premature neuronal differentiation and neuronal cell death was observed (Hunnicutt et al., 2012). While this led to a reduction in dorsal root ganglia size, an increase in axon branching from cell bodies was also observed (Hunnicutt et al., 2012). A second group uncovered increased, aberrant branching of dorsal root ganglia neurons and misguided axons after Elp1 knockdown in chick trunk neural crest cells (Abashidze et al., 2014). Our findings complement these studies in that at least some placode cells differentiate into neurons and migrate to the ganglionic anlage. Further, we noted axons with abnormal branching, especially at E2.5 (HH15-17) and in the maxillomandibular branch at E3.5 (HH19-20).

While our data revealed that knockdown of Elp1 in chick trigeminal placode cells disrupts initial trigeminal ganglion formation, this is in contrast to findings in the mouse using a conditional knockout approach in which *Elp1* is deleted from neural crest cells (Leonard et al., 2022). Subsequent development in the mouse showed disorganization and straying of trigeminal sensory axons with defects in axonal pathfinding and target innervation, also noted in our data. While some differences could be due to species variability, it is likely that many are due direct effects on placode-derived neurons instead of neural crest cells. Noting earlier phenotypic changes is logical because perturbations are occurring to the first population of trigeminal sensory neurons that are present, given that neural crest cells differentiate into neurons later than placode cells (D’Amico-Martel & Noden, 1983, Hamburger, 1961, Méndez-Maldonado et al., 2020).

Additionally, the ophthalmic branch frontal nerve initially extends axons around the eye in the Elp1 conditional knockout (but less than seen in control littermates), followed by a retraction of these axons (Leonard et al., 2022). This demonstrates that at least some axons can initially reach their destinations. Conversely, the ophthalmic medial and lateral nasal nerves fail to form in this mouse, whereas in our data, innervation, albeit reduced, is observed by all these nerves. The unique phenotypes among branches and between the two cell types investigated suggest a potential difference in origin of these neuronal derivatives in formation of the different trigeminal ganglion branches. Moreover, previous studies have reported increases or decreases in axon/neurite branching after Elp1 knockdown depending upon context (Abashidze et al., 2014; Hunnicutt et al., 2012; Jackson et al., 2014; Ohlen et al., 2017), which could be due to timepoints examined or the identity and environment of specific neurons. Later chick developmental stages comparable to these mouse studies will be beneficial to investigate to distinguish possible context-dependent differences in Elp1 function between neural crest and placode cell derivatives of the trigeminal ganglion.

### Potential mechanisms underlying trigeminal ganglion phenotypes observed after Elp1 knockdown

The deficits seen in neuron projections upon Elp1 knockdown in chick trigeminal placode cells suggest defects in axon pathfinding and guidance. Receptors for neurotrophins such as Tropomyosin receptor kinases (Trks) are important for stimulation of intracellular signaling cascades associated with growth and survival of neuron populations. Elp1 is reported to modulate TrkA/NGF retrograde signaling by regulating the phosphorylation of TrkA receptors in signaling endosomes (Li et al., 2020). One hypothesis is that neurons cannot receive NGF (i.e., insufficient TrkA is present to bind NGF), leading to their death. However, findings also support defects prior to NGF binding, including a reduction in TrkA immunoreactivity and neuron number in the Elp1 conditional knockout mouse, suggesting additional explanations for the observed phenotypes (Leonard et al., 2022). Moreover, studies in the mouse trigeminal ganglion identified a primarily neural crest origin for TrkA neurons, while TrkB and TrkC neurons arise from placode cells. If this relationship is conserved, it is likely TrkB and/or TrkC neurons are being affected when Elp1 is depleted from chick trigeminal placode cells. Immunohistochemistry on MO-treated embryonic trigeminal ganglion tissue using antibodies for TrkA, B, and C will address potential conservation of trigeminal ganglion development between aves and mammals.

Other mechanisms and pathways that may be altered upon Elp1 reduction, such as those affecting cytoskeletal regulation, cell adhesion, and vesicular trafficking, have been previously reported (Abashidze et al., 2014; Goffena et al., 2018; Johansen et al., 2008, Li et el., 2020) and are consistent with results from our functional studies. Defects in axon outgrowth could arise from cytoskeletal dysfunction, and the distribution of neural crest cells and placode-derived neurons within the trigeminal ganglion could arise due to defects in cell adhesion. Investigating possible impacted pathways will be helpful to identify mechanisms underlying Elp1-mediated trigeminal ganglion development.

Overall, our data provides new and important insights into Elp1 function in the proper development of not only placode-derived neurons, but also the trigeminal ganglion as a whole. Reducing Elp1 levels in chick trigeminal placode cells negatively impacted trigeminal ganglion development. In contrast to the loss of Elp1 in the mouse neural crest cell lineage, earlier defects in trigeminal ganglion development were observed, underscoring the known developmental timing differences in the neuronal contributions of each cell population to this ganglion. Our findings are the first to characterize the role of Elp1 in trigeminal placode-derived neurons and add to the growing literature regarding the importance of Elp1 during normal trigeminal ganglion development and how abrogation of Elp1 function leads to trigeminal nerve deficits in FD.

## EXPERIMENTAL PROCEDURES

### Chicken embryos

Fertilized chicken eggs (*Gallus gallus*) were obtained from the University of Maryland (College Park, MD) and/or outside vendors (e.g., Centurion, GA). The eggs were incubated at 37°C in a humidified incubator to desired developmental timepoints. After incubation, eggs were windowed to gain access to the embryo. India ink (Pelikan) diluted in Ringer’s solution (123mM NaCl, 1.5mM CaCl_2_, 4.96mM KCl, 0.81mM Na_2_HPO_4_, 0.147mM KH_2_PO_4_, pH 7.4) was injected under the embryo using a 1 mL syringe with an 18-gauge needle to provide contrast when viewing the embryo under a dissecting microscope. Staging of embryos was then conducted according to the Hamburger-Hamilton staging guide (Hamburger & Hamilton, 1992).

### Morpholinos

A 3’ lissamine-tagged Elp1 translation-blocking MO (5’–CAGCAGCCGCAGATTCCTCATGG– 3’) and Elp1 splice-blocking MO (5’–CCTGACAGACGCCTCACCGACTG–3’) were designed according to the manufacturer’s criteria (GeneTools, LLC). These MOs were mixed and used at a 1:1 ratio (2 mM total concentration). A standard scrambled Control MO (5’– CCTCTTACCTCAGTTACAATTTATA–3’) prepared by the manufacturer served as a control. These MOs were used at a concentration of 2mM.

The specificity of all MO sequences was verified using the NCBI Nucleotide BLAST tool. According to directions from the manufacturer, the inverse complement of the MO sequence was compared with the chicken (*Gallus gallus*) transcriptome to ensure homology was specific to desired targets, thereby mitigating the possibility of off-target effects.

### 2.4 In ovo ectodermal electroporation

Electroporation of the embryo ectoderm to target forming placode cells was performed as previously described (Shah et al., 2017). Briefly, a 27 1/2-gauge needle was used to make a hole above the embryo head, in line with the ectodermal edge, just outside the area opaca. A 1 mm x 0.75 mm 4-inch glass needle was inserted through the vitelline membrane into the space where the embryo is housed. MOs were then overlaid on top of the ectoderm at E1.5 (HH10-11; prior to placode cell delamination) to target forming trigeminal placode cells and their neuronal derivatives. Platinum electrodes (0.5 mm thick) were placed with the positive electrode on top of the chick embryo and the negative electrode beneath. The lissamine tag imparts the MO with a slightly positive charge. With the negative electrode under the embryo, the MO is thus pulled down into the ectodermal cells. Three pulses of 9V lasting 50 milliseconds with intervals of 200 milliseconds were then applied using an electroporator (California Institute of Technology). Eggs were then re-sealed using packing tape followed by parafilm and re-incubated for the desired amount of time depending on the experiment.

### Embryo collection, embedding, and sectioning

Embryos were dissected off the yolk at specific stages using the Hamburger and Hamilton staging guide then rinsed in Ringer’s solution and extra tissue surrounding the embryo was trimmed away. Embryos were then fixed in 4% paraformaldehyde (PFA) via submersion and gentle agitation at 4°C overnight (or 45 minutes at room temperature for wildtype immunohistochemistry). After fixation, embryos were permeabilized and washed three times in 1X phosphate-buffered saline (PBS) with 0.1% Triton X-100 (PBS-Tx) for 10 minutes each. Embryos used for whole-mount immunohistochemistry were stored in 1X PBS until ready for processing.

All other embryos were put through a sucrose gradient, 5% sucrose (w/v) in 1X PBS at room temperature until embryos sank (∼5-10 minutes), followed by 15% sucrose (w/v) in PBS at 4°C overnight. Embryos were then equilibrated into gelatin by submerging them in 7.5% gelatin (made in 15% sucrose) at 37°C for 8 hours and then transitioning into 20% gelatin (made in 1X PBS) at 37°C overnight. After these equilibration steps, embryos were positioned into molds containing 20% gelatin. The gelatin was allowed to solidify and then the entire mold was set on ice for 10 minutes. Next, the molds containing the embedded embryos were frozen in liquid nitrogen vapor and each embedded embryo was stored at -80°C until further processing. Embryos were sectioned using a cryostat (Leica) at 12-14μm, and sections were collected on Superfrost Plus charged slides (VWR, 48311-703). Sectioned tissue was used immediately or stored at -20°C until processed for immunohistochemistry.

### Immunohistochemistry and tissue clearing

#### Tissue sections

Slides containing sectioned tissue were either used immediately or removed from storage at -20°C and brought to room temperature. All slides were degelatinized in 1X PBS at 42°C for 20 minutes. After degelatinization, slides were placed in PBS-Tx for 30 minutes. Tissue was permeabilized in 1X PBS/0.5% TritonX-100 for 5 minutes and then blocked in 10% heat-treated sheep serum (HTSS) diluted in PBS-Tx for 1 hour. Primary antibodies were diluted in PBS-Tx + 5% HTSS, according to individual antibody dilutions (Table 1) and incubated in a humidified chamber overnight at 4°C. Slides were then washed four times for 30 minutes each at room temperature in PBS-Tx to remove any unbound primary antibodies. Sections were then incubated with secondary antibodies, diluted at 1:500 in PBS-Tx + 5% HTSS, in a humidified chamber for 1 hour at room temperature. Slides were then washed four times for 30 minutes each at room temperature in PBS- Tx to remove unbound secondary antibodies, followed by two rinses in 1X PBS for 5 minutes each, all at room temperature. Coverslips were mounted with DAPI Fluoromount-G Mounting Medium (Southern Biotech, 0100-20) and dried in the dark at room temperature overnight prior to imaging. After drying overnight, slides were stored at 4°C when not being imaged.

**Table 1:** Antibodies used for immunohistochemistry (IHC) and immunoblotting (WB).

| Antibody | Source | Catalog Number | Use/Dilution |
| --- | --- | --- | --- |
| Elp1 (IKBKAP): mouse monoclonal) | Sigma | WH0008518M3 | IHC (1:50) |
| Tubb3 (β-tubulin III): mouse monoclonal | Abcam | ab78078 | IHC (1:500 for sections;<br>1:250 for whole-mount) |
| Sox10: rabbit polyclonal | GeneTex | GTX128374 | IHC (1:500) |
| E-Cadherin: mouse monoclonal | BD Transduction<br>Laboratories | 610182 | IHC (1:500) |
| Elp1 (IKBKAP/IKAP): rabbit polyclonal | LifeSpan BioSciences | LS-B571 | WB (1:10,000) |
| Beta-actin: mouse monoclonal | Santa Cruz | sc-47778 | WB (1:10,000) |
| HRP-conjugated anti-rabbit IgG | Rockland | 611-1302 | WB (1:15,000) |
| HRP-conjugated anti-mouse IgG1 | Jackson<br>ImmunoResearch | 115-035-205 | WB (1:15,000) |

### Whole-mount

Embryos set aside for whole-mount immunohistochemistry (stored in 1X PBS at 4°C) were blocked in PBS-Tx + 10% HTSS for up to two hours at room temperature with gentle agitation. Embryos were then incubated with primary antibody (Table 1) diluted in PBS-Tx + 5% HTSS for up to 2 days at 4°C with gentle agitation. Embryos were then washed four times for 30 minutes each in PBS-Tx at room temperature, then incubated with secondary antibody diluted at 1:500 in PBS-Tx + 5% HTSS overnight at 4°C with gentle agitation. After secondary incubation, embryos were washed four times for 30 minutes each in PBS-Tx, followed by two washes with 1X PBS for 20 minutes each, all at room temperature. E2.5 (HH15-17) embryos were imaged at this step, while E3.5 (HH19-20) embryos were cleared before imaging, as described below.

#### 2.6.3 Fructose and urea solution (FRUIT) clearing

After whole-mount immunohistochemistry, embryos were cleared using the FRUIT clearing method to better visualize the forming trigeminal ganglion (Hou et al., 2015). Embryos were moved through a series of FRUIT buffers containing 8M urea (Sigma, U5378), 0.5% (v/v) α- thioglycerol (Thermo Fisher, T090525G), with increasing concentrations of fructose (Sigma, F3510). Embryos were incubated in 35% FRUIT for 6 hours, 40% FRUIT overnight, 60% FRUIT for 8 hours, followed by 80% FRUIT overnight. All incubations were performed at room temperature with gentle rocking. Embryos were stored and imaged in 80% FRUIT buffer. Embryos were not stored in 80% FRUIT for more than 2 days prior to imaging due to crystallization occurring if embryos are left for too long.

### Confocal imaging

All imaging was performed on a Zeiss LSM 800 confocal microscope. Tissue sections were imaged using 5X, 10X, or 20X air objectives, or the 63X oil objective. Embryos processed for whole-mount immunohistochemistry were imaged in either 1X PBS (Elp1 MO, E2.5 (HH15-17) or 80% FRUIT. Z-stack images were taken at 5 µm intervals using 5X and 10X air objectives. For all applications, laser power, gain, offset, and digital zoom remained the same for Contralateral vs. MO-treated tissue. CZI files were processed using Zen software (Blue edition 2.0, Zeiss) while Z-stack CZI files were processed in ImageJ using the Z-project function (Hyperstack mode) to create maximum intensity projections.

### Measurements and Statistical Analysis

Measurements were conducted using the open-source image processing program FIJI (Schindelin et al., 2012), which is based on ImageJ software (Collins, 2007). Measurements were performed on Tubb3-labeled maximum intensity Z-projections described above. To measure area, the boundary of the trigeminal ganglion was determined by Tubb3 staining and outlined using the freehand sections tool then measured. Ectodermal Tubb3 staining was cropped out to better visualize the trigeminal ganglion, as it merges with maxillomandibular branching of the trigeminal ganglion when creating maximum intensity Z-projections.

Similarly, measurements of the area occupied by axon projections from the ophthalmic branch that innervate the eye were obtained by outlining the nerve innervation starting at the point at which additional branches break away from the main ophthalmic branch. Contralateral and MO-treated trigeminal ganglia were compared using paired t-tests in Graphpad Prism, with *p* values less than or equal to 0.05 were considered significant.

### Tissue collection

Embryos electroporated with Elp1 MO or the standard Control MO (described above in 2.2), were collected 48 hours post-electroporation. Successfully electroporated trigeminal ganglia were dissected in 1X PBS under a Zeiss SteREO Discovery V8 Pentafluor fluorescent microscope to visualize MO-positive fluorescent tissue and then pooled for immunoblotting (n = 18 Control MO, n = 21 Elp1 MOs). The tissue was pelleted by centrifuging at 500 x g for 5 minutes at 4°C and excess buffer was removed, followed by flash-freezing of tissue in liquid nitrogen, and storage at -80°C until ready for immunoblotting.

### Immunoblotting

Tissue pellets were thawed on ice and lysed in lysis buffer (50mM Tris pH 8.0, 150mM NaCl, 1% IGEPAL CA-630) supplemented with 1X cOmplete protease inhibitor cocktail (Roche, 04693124001) and 1mM PMSF (Roche, 54561623) for 30 minutes on ice, gently vortexing every 10 minutes. Samples were then centrifuged at max g for 15 minutes at 4°C and the solubilized protein fraction was collected. Protein concentration was determined using a Bradford assay (Thermo, 1857210) and equivalent amounts of protein per sample were boiled in 1X Laemmli sample buffer at 99°C for 5 minutes, followed by cooling to room temperature.

Prepared samples were loaded onto a 7.5% (Elp1) Sodium dodecyl sulfate–polyacrylamide gel electrophoresis (SDS-PAGE) (Biorad 4561033, 4561024 respectively) and separated by electrophoresis at 70V for approximately 2.5 hours in 1X running buffer (diluted from 10X running buffer (25mM Tris base, 1.92M Glycine, 3.5mM SDS)). To transfer proteins from the gel, 0.2 µm PVDF membrane (Thermo Fisher, 88520) was first equilibrated for 30 seconds in methanol followed by a 20-minute equilibration in 1X transfer buffer (100mL 10X running buffer, 200 mL methanol, 700 mL ddH_2_O). After protein separation, the SDS-PAGE gel was incubated in 1X transfer buffer for 20 minutes for equilibration. Proteins were then transferred to PVDF membrane using the Mini Trans-Blot ® Cell (Bio-Rad) wet transfer system running at 70V for 2 hours at 4°C. Membranes were incubated in a blocking solution (1X PBS + 0.1% Tween-20 (PTW) + 5% milk) for 45 minutes at room temperature followed by primary antibody incubation overnight at 4°C, also diluted in blocking solution (Table 1). After primary antibody incubation, membranes were washed in PTW three times for 10 minutes each. Secondary antibodies (Table 1) were diluted in blocking solution for one hour at room temperature followed by three additional washes in PTW for 10 minutes each. Membranes were incubated with chemiluminescent substrates (Supersignal West Pico PLUS (Thermo Fisher, 34580) and/or Supersignal West Femto (Thermo Fisher, 34095)), and developed using a ChemiDoc XRS system (Bio-Rad). Blots were then stripped using Restore Plus Western Blot Stripping Buffer (Thermo Fisher, 46430) for 15 minutes, rinsed two times in PTW, and re-blocked for 45 minutes in blocking solution. Membranes were then re-probed using β-actin (Table 1) as a loading control antibody following the above procedures. Immunoblot analysis was performed using Image Lab software (Bio-Rad) to determine band size and volume. Relative protein levels were calculated by normalizing to loading control band volumes. Differences in amounts of protein were determined by comparing normalized ratios between Control MO- and Elp1 MO-treated samples, with Control MO-treated samples set to one.

## ACKNOWLEDGMENTS

This work was supported by NIH R01DE024217 and R03HD108480 (L.A.T.) and the Ann G. Wylie Dissertation Fellowship from the University of Maryland Graduate School (M.A.H.).

